# Identification of strain-specific cues that regulate biofilm formation in *Bacteroides thetaiotaomicron*

**DOI:** 10.1101/2024.12.20.629428

**Authors:** Robert W.P. Glowacki, Morgan J. Engelhart, Jessica M. Till, Anagha Kadam, Ina Nemet, Naseer Sangwan, Philip P. Ahern

## Abstract

Members of the gut microbiome encounter a barrage of host- and microbe-derived microbiocidal factors that must be overcome to maintain fitness in the intestine. The long-term stability of many gut microbiome strains within the microbiome suggests the existence of strain-specific strategies that have evolved to foster resilience to such insults. Despite this, little is known about the mechanisms that mediate this resistance. Biofilm formation represents one commonly employed defense strategy against stressors like those found in the intestine. Here, we demonstrate strain-level variation in the capacity of the gut symbiont *Bacteroides thetaiotaomicron* to form biofilms. Despite the potent induction of biofilm formation by purified bile in most strains, we show that the specific bile acid species driving biofilm formation differ among strains, and uncover that a secondary bile-acid, lithocholic acid, and its conjugated forms, potently induce biofilm formation in a strain-specific manner. Additionally, we found that the short-chain fatty acid, acetic acid, could suppress biofilm formation. Thus, our data defines the molecular components of bile that promote biofilm formation in *B. thetaiotaomicron* and reveals that distinct molecular cues trigger the induction or inhibition of this process. Moreover, we uncover strain-level variation in these responses, thus identifying that both shared and strain-specific determinants govern biofilm formation in this species.

**Importance:** In order to thrive within the intestine, it is imperative that gut microbes resist the multitude of insults derived from the host immune system and other microbiome members. As such, they have evolved strategies that ensure their survival within the intestine. We investigated one such strategy, biofilm formation, in *Bacteroides thetaiotaomicron*, a common member of the human microbiome. We uncovered significant variation in natural biofilm formation in the absence of an overt stimulus among different *Bacteroides thetaiotaomicron* strains, and revealed that different strains adopted a biofilm lifestyle in response to distinct molecular stimuli. Thus our studies provide novel insights into factors mediating gut symbiont resiliency, revealing strain-specific and shared strategies in these responses. Collectively, our findings underscore the prevalence of strain-level differences that should be factored into our understanding of gut microbiome functions.

## Introduction

*Bacteroides thetaiotaomicron* is a prominent member of the microbiome present in the human gastrointestinal tract (1). Given its abundance in human populations and genetic tools for its manipulation, the mechanisms through which it competes for nutrients in the intestine and imparts beneficial features to the host have been studied in detail (2–12). Although some mechanisms that foster intestinal resilience in *B. theta* have been uncovered, there remains a void in our understanding of how *B. thetaiotaomicron* can persist in a dense, competitive environment in the face of ecological challenges such as host-produced anti-microbial peptides (13), and reactive oxygen species (14). One mechanism that provides resistance to a variety of stressors in microorganisms is biofilm formation (15, 16). Biofilms are aggregates of microbial cells that are embedded in extracellular polymeric substances that can shield the microbes from harmful agents, and as such, represent a lifestlye that allows microbes to survive and thrive in harsh environments (16, 17). Although biofilm formation has not been extensively studied in symbiotic members of the microbiome, recently, it has been uncovered that the type strain of *B. thetaiotaomicron* (*Bt*), *Bt*-VPI-5482, forms robust biofilms in response to purified bile, but not in the absence of an overt stimulus (so-called “natural” or “inherent” biofilm-forming capacity) (18). Seminal studies of this process have uncovered the genetic circuitry that regulate aspects of the biofilm formation in *Bt*-VPI-5482, revealing pathways that mediate repression of inherent biofilm-forming capacity (19, 20), or promotion of biofilm formation in response to bile (18, 21) highlighting that biofilm formation appears to be critically tuned to environmental cues.

Despite the clear impact of bile on biofilm formation, several important questions have remained unanswered. Bile is a mixture of many distinct compounds including retinol (22), bile acids (23), cholesterol (24), bilirubin (25), and more. However, the precise molecular components of bile that promote biofilm formation in *B. theta* have not been defined. Moreover, as our appreciation of the prevalence and importance of strain-level variation in gut microbes with respect to microbial fitness and beneficial host effects grows (26–29), it is important that such characteristics are determined in multiple strains to determine if they represent broad features of a species or strain-specific pathways that promote strain fitness in a given context. In addition, the existence of inhibitors of biofilm formation in *B. theta* is unknown, creating a gap in our understanding of the environmental signals that regulate the formation of biofilms. Given that biofilm formation can contribute to microbial pathogenesis, and that *B. thetaiotaomicron* also harbors pathogenic potential in the intestine (30) (31–34), understanding how biofilm responses are initiated and repressed, and how this varies depending on the nature of the strain of interest may be important for strategies that aim to modulate the function of the microbiome for therapeutic purposes.

Here we investigated a panel of human gut-derived *B. thetaiotaomicron* strains to examine their intrinsic biofilm forming abilities, representing several strains whose biofilm forming capacity was unknown. We identified a new set of strains with the ability to form robust biofilms in the absence of bile as an inducing molecule, thus uncovering extensive strain-level variation in this response. One of these strains, *B. thetaiotaomicron* VPI-5951 (henceforth referred to as *Bt*-5951) was further investigated. By contrast with *Bt*-VPI-5482, we determined that biofilm formation by *Bt*-5951 was not substantially bile-responsive but that lithocholic acid (LCA), a secondary bile acid, and its taurine and glycine-conjugated forms were able to induce *Bt-*5951 to form robust biofilms substantially above that of intrinsic levels, thus revealing variation with respect to the stimuli that induce or expand biofilm formation. Further, we show that LCA glyco and tauro conjugates, glycolithocholic acid (GLCA) and taurolithocholic acid (TLCA), also act as a biofilm inducer in several strains, suggesting these bile acids represent a broad biofilm inducing signal. Moreover, we find that for strain *Bt*-5951, none of the other single bile acid substrates tested is able to support biofilm formation. Additionally, we show that acetic aicd, a microbial-derived short-chain fatty acid (SCFA), suppresses the intrinsic biofilm forming capability of strains *Bt-5951* and *Bt*-0940-1. Thus, our data (i) identifies strains of *B. theta* that had not previously been known to have inherent biofilm-forming capacity, (ii) define key physiologically relevant molecules that govern the formation and inhibition of biofilms in *B.* theta, and (iii) uncover strain-selectivty in the cues that regulate this process, thus providing novel insights into strain-level variation in *B. theta*. Therefore, our approach expands our understanding of the gut-relevant molecules that coordinate biofilm formation in *B. thetaiotaomicron* and bolsters the utility in studying non-type strains to study the physiology of gut *Bacteroides*.

## Results

### Inherent biofilm formation ability differs among *B. thetaiotaomicron* strains

To identify *B. thetaiotaomicron* isolates with intrinsic biofilm formation capacity, 22 human-derived *B. thetaiotaomicron* isolates were grown on tryptone-yeast extract-glucose (TYG) medium without the presence of known *Bacteroides-*specific biofilm inducers (18, 35, 36). While the majority of *B. thetaiotaomicron* strains failed to form potent biofilms in TYG media across multiple experiments, (**Fig 1A**), three strains, *Bt-*5951, *Bt*-0940-1, and *Bt*-3443, all formed robust biofilms compared to the type strain, VPI-5482 (*Bt*-VPI-5482) (**Fig 1B, Fig S1A-C**). To determine if this was TYG-media specific or if it represents an intrinsic feature in these strains, we grew these strains in Brain-Heart Infusion-Supplemented (BHIS) media and found similar biofilm-forming pattern among strains (**Fig 1C**), albeit with differing magnitudes of biofilm formation than that seen in TYG. We next selected strain *Bt-5951* as a representative strain for further study due to its intrinsic biofilm forming capabilities and its potential for future followup studies due to its genetic tractability, while strain *Bt*-0940-1 and *Bt*-3443 have proved largely recalcitrant to genetic modification to date. Next, these intrinsic biofilms formed by strain *Bt-5951* were treated with DNase, RNase, or Proteinase K, revealing that all three treatments substantially impaired biofilm formation of TYG-grown *Bt-*5951 (**Fig D**), in keeping with the known role for their targets in the establishment of a biofilm (37). With all three enzymes, residual biofilm was seen, suggesting other factors such as carbohydrates or lipids could be incorporated into these intrinsic biofilms of *Bt-*5951 as shown in other species of bacteria (17, 38). Additionally, supplementation of the media with the nonionic surfactant Tween 80 was able to completely inhibit biofilm formation (**Fig 1D**) similar to previous reports of nonionic surfactants inhibiting biofilm or cell attachment, suggesting this is a *bona fide* biofilm structure (39).

**Figure 1.**
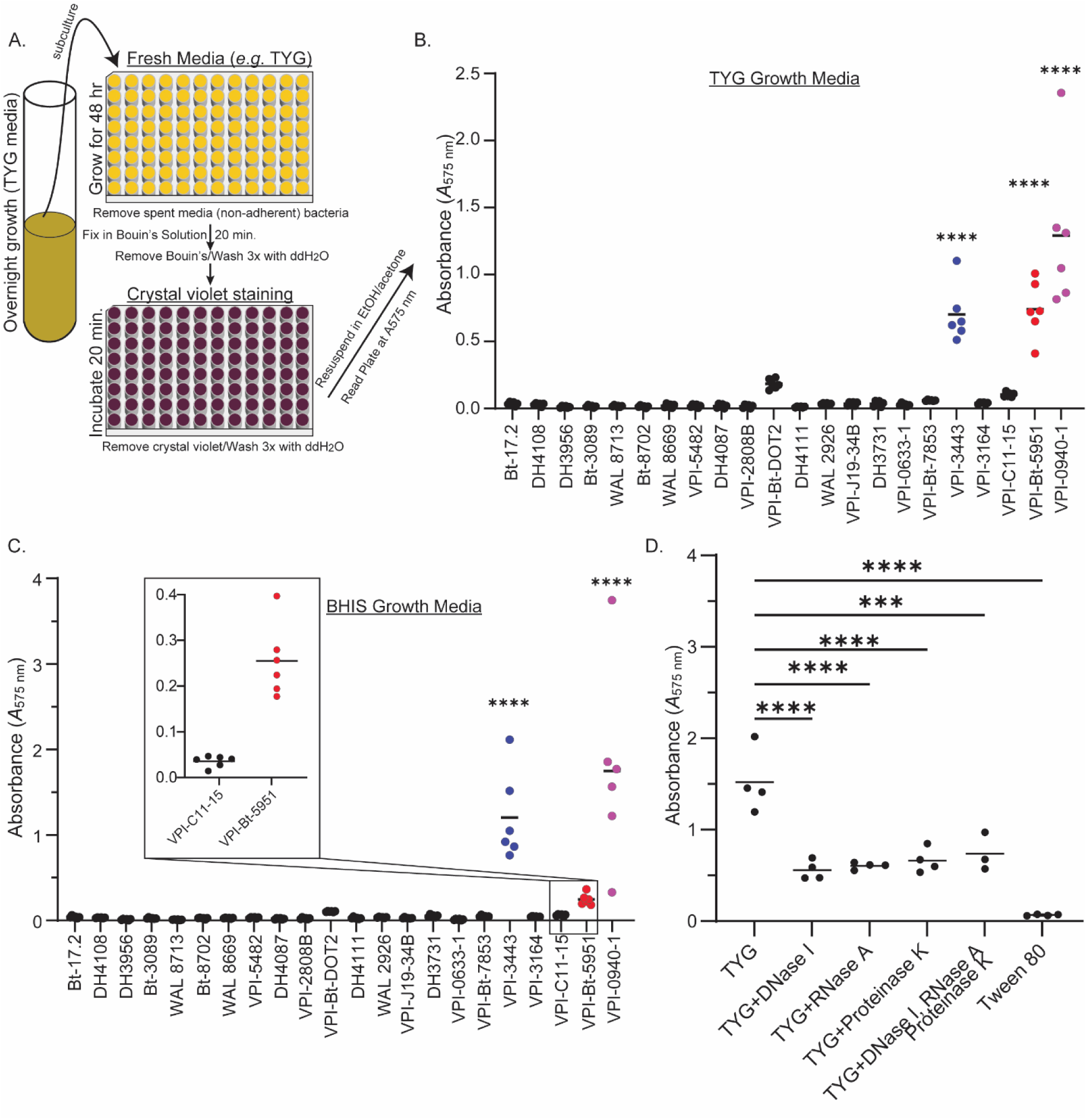
Inherent biofilm formation ability differs among *B. thetaiotaomicron* strains. (A) Schematic of the crystal violet-based assay for biofilm formation used throughout this study (B-C) 22 isolates of *Bacteroides thetaiotaomicron* were grown in the indicated growth medium and their biofilm forming capacity was assessed using a crystal violet-based biofilm assay, schematic in (A). Biofilm formation following growth in (B) TYG or (C) BHIS was quantified by assessment of absorbance at 575 nm (*A*575) following crystal violet staining and is shown for each of the indicated strains at 48 hours post-inoculation. Inset in (C) shows the magnitude of strain *Bt*-5951 biofilm formation. (D) The impact of DNase, RNase, and Proteinase K treatments on biofilm formation in strain *B. theta*-5951 was assessed at 48 hours. Data points show individual technical replicates, and bars represent the mean. Graphs are representative of 4 and 3 independent experiments in (B, C) respectively. Statistical significance was determined using a one-way ANOVA with Dunnet’s multiple comparisons to strain *Bt*-*VPI*-5482 in (A) and (B), and to TYG in (C). *p* values of <0.05 were considered statistically significant *\*\*\*, p*<0.001 and ****, *p*<0.0001.

### Lithocholic acid is a potent biofilm inducer in *B. thetaiotaomicron* strain 5951

Given the striking capacity of bile to induce biofilm formation in *Bt*-VPI-5482 (18), we sought to address whether biofilm formation capacity in response to bile was a generalizable feature of the strains we tested. We focused on the following strains: the type strain *Bt*-VPI-5482 as a positive control and two of the intrinsic biofilm formers *Bt-*5951 and *Bt*-0940-1. The type strain, *Bt*-VPI-5482, showed substantial induction in biofilm formation in a dose-dependent manner (**Fig 2A**). Similarly, strain *Bt-*0940-1 showed a substantial increase in biofilm formation in response to bile. Furthermore, the bile-induced *Bt*-VPI-5482 and *Bt*-0940-1 biofilms were treated with DNase, RNase, and Proteinase K and we observed significant decreases following treatment with DNase and Proteinase K, as previously reported for bile-induced biofims (**Fig S2A,B**), further validating our findings. Strain *Bt-*5951, unlike other tested strains, did not increase biofilm production with bile supplementation of up to 2% w/v (**Fig 2A**). These data suggest that strain *Bt*-5951 does not form biofilms in response to bile, contrasting with other strains, and/or it requires higher levels of the same stimulus than is present in bile than the other strains tested to stimulate biofilm formation.

**Figure 2.**
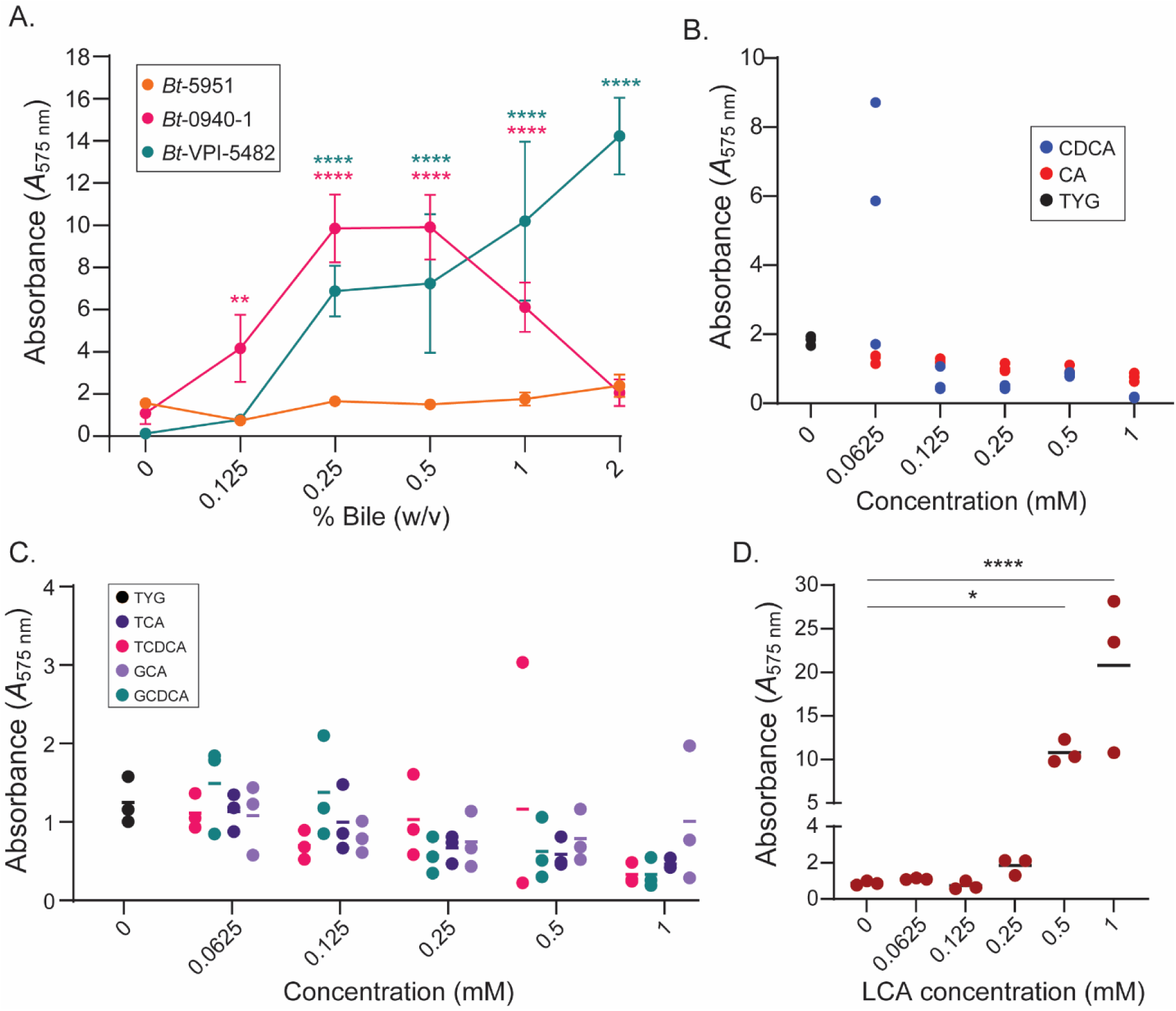
Only the secondary bile acid lithocholate elicits biofilm formation in strain *Bt*-5951. (A) Bile-induced biofilm formation at 48 hours as assayed by crystal violet biofilm staining in *Bt* strains 5482, 5951, and 0940-1. Data points show the mean of an average of 4 technical replicates, error bars show standard deviation. (B) Biofilm formation dose response at 48 hours for strain *Bt*-5951 grown in TYG media containing primary bile acids cholic acid (CA) or chenodeoxycholic acid (CDCA). (C) Biofilm formation dose response at 48 hours for strain *Bt*-5951 grown in TYG media containing taurine and glycine conjugated bile acids taurocholic acid (TCA), taurochenodeoxycholic acid (TCDCA), glycocholic acid (GCA), or glycochenodeoxycholic acid (GCDCA). (D) Biofilm formation dose response at 48 hours for strain *Bt*-5951 grown in TYG media containing lithocholic acid (LCA) Graphs are representative of 3 independent experiments in (A), while (B-D) the dose response is representative of 2 independent experiments, (data for 0.5 mM for all bile acids is representative of at least 3 independent experiments). Statistical analysis in (A) was performed using a two-way ANOVA with Dunnet’s multiple comparisons, the mean of each data point within each strain was compared to the 0 mM point. Statistical analysis in (D) was performed using one-way ANOVA with Dunnet’s multiple comparisons to the mean of TYG without LCA (0 mM). *p* values of <0.05 were considered statistically significant, *, *p*<0.05, ****, *p*<0.0001. Significance astrisks in (A) are color-coded to match the strain.

As bile is a complex mixture of cholesterol and lipid molecules, bile acids (both host primary and microbiota-generated secondary), vitamins, bilirubin, and other trace compounds (22–25, 40), we reasoned that strain *Bt*-5951 may respond to a bile acid species that was either not present or under-represented in the bile mixture we used to induce biofilm formation in *Bt*-VPI-5482. To identify components of bile that have the capacity to induce biofilm formation we further tested the capacity of individual bile acids and their glycine and taurine conjugates (i.e. bile acid salts) that are commonly found in the intestines through the actions of the liver or the gut microbiome (41, 42) to elicit a biofilm response in strain *Bt*-5951. First, we tested two primary, host derived, bile acids chenodeoxycholic acid (CDCA) and cholic acid (CA), and observed no change in the amount of biofilm formed when compared to control media. (**Fig 2B**). Next, we tested glycine and taurine-conjugated forms of CDCA and CA, and again we did not observe a statistically significant increase in biofilm formation (**Fig 2C**). Finally, we tested secondary bile acids produced by microbial modification of host-derived bile acids and observed that lithocholic acid (LCA) induced a nearly 400% increase in biofilm formation over that seen for an intrinsic biofilm at a concentration of 1 mM for *Bt*-5951 (**Fig 2D**). This effect was specific to LCA as other secondary bile acids tested, including deoxycholic (DCA), taurodeoxycholic acid (TDCA), ursodeoxycholic acid (UDCA), and hyodeoxycholic acid (HDCA) did not elicit signficant increases in biofilm formation (**Fig S2C**), although we cannot exclude that additional microbiome-produced bile species could mediate similar effects or that they are important in other contexts. Due to its low solubility, LCA precipitates from the solution over time at concentrations of 1 mM and higher. However, HDCA and UDCA also form precipitates and do not stimulate biofilm formation (**Fig S2C**), demonstrating that precipitation alone is not responsible for the observed biofilm induction. Moreover, at 0.5 mM we did not observe precipitation of LCA, yet saw large increases in biofilm formation, further reinforcing that LCA itself rather than the presence of precipitate is responsible. Furthermore, LCA alone controls without bacteria present also precipitated at 1 mM but had low background compared to when bacteria are present (**Fig S2D**), confirming that the precipitated LCA itself did not contribute substantially to the signal. Finally, the solvents used to prepare these secondary bile acid stocks, DMSO or ethanol (EtOH), did not significantly impact biofilm formation when added to TYG, reflecting that LCA is sufficient to cause biofilm formation irrespective of the specific solvent present. (**Fig S2E**).

Next we tested conjugated forms of LCA, taurine-LCA (TLCA), and glycine-LCA (GLCA), as well as its iso-, oxo-, and alloiso-forms. We observed that both TLCA and GLCA caused a substantial increase in biofilm formation in *Bt-*5951, even greater than those observed with LCA (**Fig 3A)**. Additionally, we observed that both iso- and alloiso-LCA produced similar levels of biofilm formation as compared to LCA (**Fig 3A**), while 3-oxo-LCA failed to cause an upregulation in biofilm formation over that of TYG (**Fig 3A**). These results clearly show that LCA, epimers of LCA, and conjugated forms of LCA promote biofilm formation in *Bt*-5951, while also demonstrating that no other bile acid species tested is capable of inducing biofilm formation in this strain.

**Figure 3.**
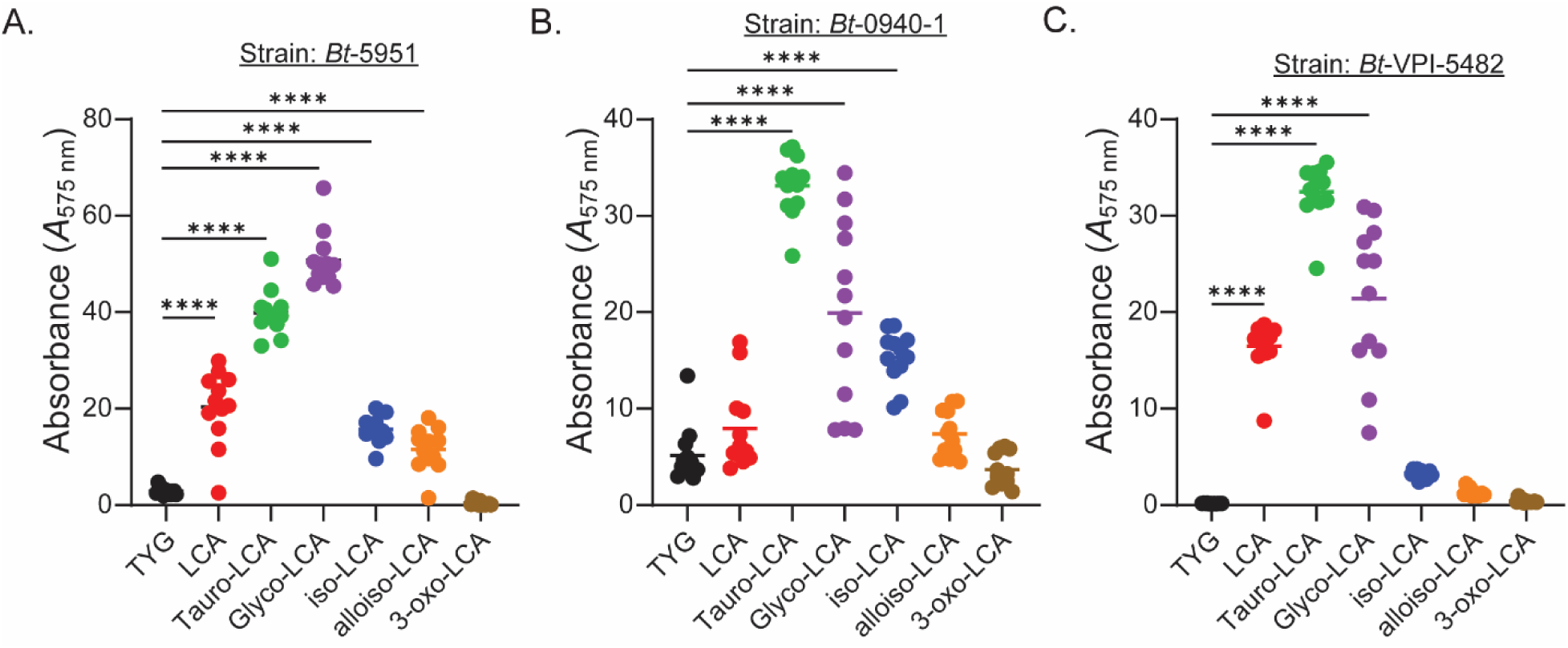
Different forms of lithocholic acid mediate biofilm formation in strains of *B. theta*. (A-D) Biofilm formation in the presence of lithocholic acid (LCA) and its epimers or conjugated forms (TYG, LCA, taurolithocholic acid, glycolithocholic acid, isolithocholic acid, alloisolithocholic acid, and 3-oxolithocholic acid) for 3 individual strains of *Bt* was assessed using a crystal violet based biofilm assay. All bile acids were tested at 0.5 mM, with the exception of isolithocholic acid which was tested at 0.25 mM due to solubility issues at higher concentrations. Strains shown are: (A) strain *Bt*-5951, (B) strain *Bt*-0940-1, (C) strain *Bt*-VPI-5482. All biofilms were measured after 48 hours of growth. Data are representative of 3 (A-C) independent experiments. Statistical significance was determined using one-way ANOVA with Dunnet’s multiple comparisons to TYG (no bile acid added). *p* values of <0.05 were considered statistically significant, ****, *p*<0.0001.

Our observations that LCA and its conjugates and isoforms could elicit biofilm formation in strain *Bt*-5951, led us to ask if this pattern was strain-specific or broadly stimulatory of biofilm formation in the species as a whole. As with *Bt-*5951, strains *Bt*-0940-1 and *Bt*-VPI-5482 showed a similar trend with no significant biofilm induction in response to most of the individual tested bile acids besides LCA (**Fig S3A,B**); it should be noted that the response to particular bile acids was somewhat inconsistent for these two strains (data for HDCA and UCDA was not consistent across experiments in *Bt*-VPI-5482 with no effects seen in some experiments and a slight increase in another; data for GCA was not consistent across experiments in *Bt*-0940-1 with no effects seen in some experiments and a slight increase in another). Strikingly, despite potent biofilm-inducing effects in *Bt*-VPI-5482, LCA did not substantially increase biofilm formation above that of intrinsic biofilm formation in *Bt-*0940-1 (**Fig 3B**), despite the strong biofilm induction driven by conjugated forms of LCA (TLCA and GLCA) as well as iso-LCA in this strain (**Fig 3B**). Furthermore, by contrast with *Bt-*5951, alloiso-LCA did not induce biofilm formation response in strain *Bt*-VPI-5482 (**Fig 3C**) compared to LCA. Thus, these individual strains respond to both shared and strain-specific cues to coordinate the formation of biofilms. In an attempt to reconcile the capacity of the identified bile acids to induce biofilm formation in strain *Bt*-5951, with the failure of purified bile to do so in this strain, we performed mass spectrometry based analysis of the bile used in our assays, as well as our LCA and TYG media (**Fig S3C,D**). Unsurprisingly, although several forms of LCA were detectable in bile at 1% w/v, even the combined concentration of all LCA species was at least an order of magnitude lower than the 0.5 mM concentration where we saw effects on biofilm formation (**Fig S3D**). This result may reflect why strain *Bt*-5951 did not increase biofilm formation in response to bile, but responded robustly to the various forms of LCA tested. Interestingly, of the 51 bile acid species that we attempted to detect, only 21 were detectable. Of those 21, the only species above 0.5 mM were, CA, GCA, TCA, TCDCA, GDCA, and TDCA which lacked potent biofilm induction capacity. Furthermore, our analysis of TYG media revealed that only trace amounts of CA are present in the nanomolar range and no bile acids that support biofilm formation were detected (not shown). Lastly, despite an advertised purity of ≥95%, an independent assessment of LCA purity in-house (not shown), showed that it was comprised of ∼50% LCA and ∼50% cholic acid (CA). However, we found no impact of cholic acid on biofilm formation for our strains tested (**Fig 2B and Fig S3A,C**) allowing us to attribute the induction to LCA.

### Short-chain fatty acids inhibit intrinsic biofilm formation

Due to our observation that bile acids, a prominent microbial-modified compound in the intestines with important physiological implications, had such a profound effect on biofilm formation, we decided to test short-chain fatty acids (SCFAs), another intestinally-abundant microbial-derived molecule of physiological relevance (43), for their role in the regulation of biofilm formation within *Bacteroides thetaiotaomicron*. We initially tested five SCFAs commonly found in the intestines and tested them at their physiological levels observed within intestinal contents from healthy individuals (43, 44). Importantly, the concentrations tested are not growth inhibitory as we tested them at or below concentrations found in “Gut Microbiota Media” (GMM), a media that supports the growth of intestinal anaerobes, including *Bacteroides* (45) and we did not observe growth defects in our assay. Strain *Bt*-5951 was largely resistant to the effects of most SCFA, but was profoundly inhibited by acetic acid (**Fig 4A**). By contrast, we observed reductions in the amount of intrinsic biofilm formed for *Bt*-0940-1 in response to most SCFA tested (sensitivity to valeric acid was not evident in all experiments), suggesting that its natural biofilm-forming capacity is exquisitely sensitive to SCFA (**Fig 4B**). We next determined the concentration of acetic acid that inhibited biofilm formation by testing the impact of a range of acetic acid concentrations on biofilm formation in strains *Bt*-0940-1 and *Bt*-5951. While both strains showed a significant reduction in biofilm formation at the higher concentrations tested, *Bt*-0940-1 was susceptible to inhibition at lower doses than strain *Bt*-5951 (**Fig 4C,D**). In order to test if reduced biofilm formation was a product of reduced viability, we performed colony forming unit (CFU) assays following incubation with our range of acetic acid concentrations and for all of the SCFAs tested. There was no significant reduction in viability for strain *Bt*-5951 following supplementation with acetic acid or other SCFAs (**Fig S4A**). Although strain *Bt*-0940-1 (**Fig S4B**) did show statistically significant reductions in growth when grown with acetic acid supplementation, these differences were likely not the driver of impaired biofilm formation as they were small in magnitude and largely indistinguishable from growth in levels of acetic acid that did not impact biofilm formation, i.e. growth was indistinguishable between 4 mM and 16 mM acetic acid, yet 4 mM acetic acid had little impact on biofilm formation.

**Figure 4.**
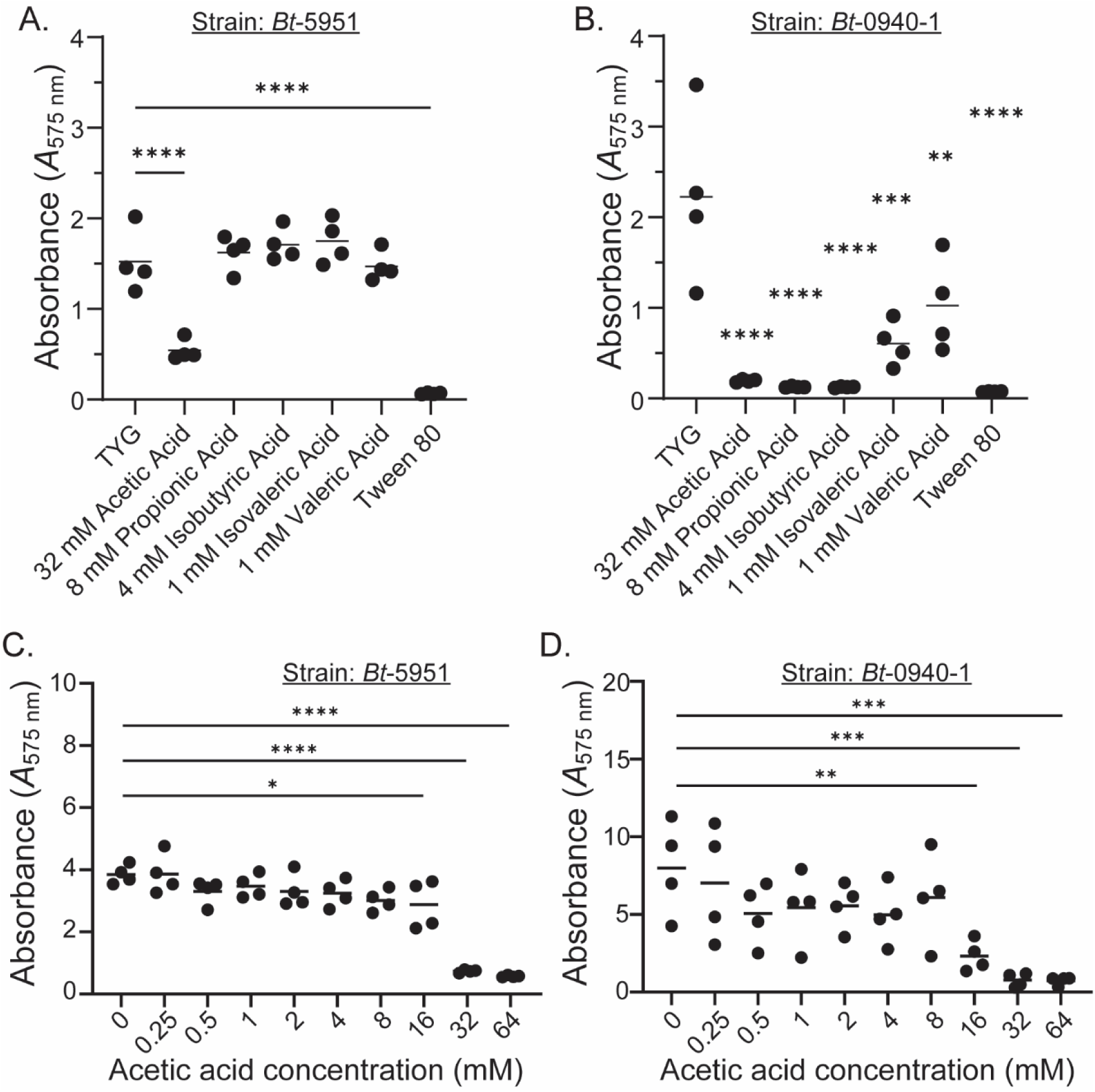
The short-chain fatty acid acetic acid inhibits the natural biofilm formation of *Bt* strains. (A-D) Biofilm formation was assessed in strains *Bt-*5951 (A and C) and *Bt*-0940-1 (B and D) using a crystal violet based biofilm assay in TYG growth media in the presence of the indicated short-chain fatty acids (SCFA) (A and B), or in a range of doses for the SCFA acetic acid (C and D). Data in (A and B) is representative of 2 independent experiments and (C and D) is representative of 3 independent experiments. Tween 80 was added at a concentration of 0.02% (v/v) into TYG media. Statistical significance was determined in (A-D) using one-way ANOVA with Dunnet’s multiple comparisons to TYG (0 mM) (no short chain fatty acid present). *p* values of <0.05 were considered statistically significant, *, *p*<0.05, **, *p*<0.01, *\*\*\*, p*<0.001 and ****, *p*<0.0001.

Together, our data define a role for intestinal bile acids and SCFAs such as acetic acid in the regulation of the biofilm-forming propensity of *B. theta* strains *in vitro*, and suggests that these two classes of molecules may operate as signaling molecules that govern biofilm formation in a strain-specific manner that influences survival in the gut against the multitude of challenges they encounter.

## Discussion

Members of the gut microbiome face myriad challenges to ensure their fitness in the intestine (46). They must thrive in the face of ever-changing nutrient sources (host diet) (47, 48), competition from other microbes who share their niche (49, 50), antibiotic exposures (51, 52), and a plethora of anti-microbial factors produced by the intestinal immune system (53–55). Despite these challenges, it is evident that many members of this complex intestinal ecosystem have evolved strategies that allow them to overcome these insults. This stability is exemplified by *B. thetaiotaomicron*, a prominent gut symbiont found in a large proportion of human microbiomes (56), that has evolved a variety of strategies that help ensure persistence in the absence of carbon sources of preference (3, 57), as well as modifications to its LPS that provide resistance to anti-microbial peptides (13). *B. thetaiotaomicron* is resistant to the stressors associated with intestinal inflammation (34), but notwithstanding some key insights into the underlying mechanisms involved (13), little is known about how this resilience is mediated. Moreover, the fact that many individuals harbor the same strains of *B. thetaiotaomicron* over extended time periods (58) suggests the existence of strain-specific strategies for survival within the intestine. While the impact of strain-level variation has been extensively studied in microbial pathogenesis (59), such variation in microbiome members has received comparatively little attention, despite evidence of its importance (26, 60–62).

The production of biofilms encased in an extracellular matrix composed of exopolysaccharides, proteins, extracellular DNA, and carbohydrates is a feature of a dense bacterial community that provides microbes a means to resist a variety of factors that could impact their survival. Although, traditionally biofilm formation studies have focused on human pathogens due to their capacity to worsen disease outcomes (63–66), little attention has been devoted to biofilm-forming abilities of resident human gut microbiome members, despite their intimate relationships within the intestine where biofilms contact the host mucosal layer, but do not interact with the host epithelium during homeostasis (67). Recent studies have uncovered biofilm formation, and signals governing biofilm initiation in several human gut commensals including *Bacteroides fragilis, B. thetaiotaomicron, Bifidobacteria,* and *Clostridial* species (18, 35, 36, 68, 69). Biofilm formation may be advantageous to these resident members of the gut microbiome as a means to withstand environmental stressors (15, 16) such as fluctuations in pH and osmolarity (70, 71), alterations to the composition of the bile acid pool (72–74), starvation due to changes in host diet (75, 76), or antimicrobial peptides (77). Here we have interrogated biofilm formation among different strains of *B. thetaiotaomicron*. Our study has uncovered significant strain-level variation in the propensity to form biofilms in the absence of any currently known inducer molecule or otherwise previously determined factor in separate media conditions, while replicating prior work showing that the type strain of *B. thetaiotaomicron* (*Bt*-VPI-5482) lacks the capacity to form a biofilm in the absence of bile (19). Of the 22 strains of *B. thetaiotaomicron* studied here, only 3 of them (strains *Bt*-0940-1, *Bt*-5951 and *Bt*-3443) displayed potent biofilm formation in the absence of bile, while strains VPI-*Bt*-DOT2 and VPI-C11-15 displayed weaker biofilm forming capacity, just above background. While we cannot exclude that the specific media types used masked natural biofilm formation in particular strains, the fact that the same phenomenon was seen using different media types suggests that natural biofilm formation is a feature of select *B. thetaiotaomicron* isolates rather than a widespread feature of the species. Moreover, although no previously described inducing molecule such as bile is present in stimulatory amounts in our media, we cannot rule out that a nutrient present in both media types is acting as an unknown trigger of biofilm formation. Thus, the distribution of biofilm-forming capacity may reflect unique strain-specific strategies that have evolved to allow strains to effectively compete for survival in the intestine, while other strains may rely more on the capacity to degrade a wide array of nutrients to maintain fitness (3, 78, 79). Given the potential for *B. thetaiotaomicron* to form abscesses under specific physiological conditions (28, 80), it is possible that this variation in biofilm formation represents a potential virulence feature in these particular contexts. Importantly, the variable distribution of this capacity has implications for studies that aim to infer/predict functions based on phylogenetic relatedness (81). Given the utility in generating accurate predictions of phenotypic outcomes based on the presence/abundance of particular microbes, defining how such phenotypes vary based on strain identity, and how representative a feature of interest is of the species as a whole is critical. Moreover, our data reinforce the the need for strain-level resolution in the prediction of community member functions.

Elegant studies have uncovered a role for bile in the induction of biofilm formation among many strains of *B. thetaiotaomicron*, including the type strain *Bt*-VPI-5482 (18–21). To our surprise, despite observing that bile induced or expanded biofilm formation in strains lacking or possessing intrinsic biofilm-forming capacity respectively, it had little to no impact on *Bt*-5951. Thus, while bile appears to be a signal for the initiation of biofilm formation among most strains tested, the effect is not uniform, and different strains may respond to distinct molecular cues to initiate this response. Although purified bile had no impact on biofilm formation in *Bt*-5951, the secondary bile acid LCA (a component of bile), its epimers, and its taurine or glycine conjugated forms expanded biofilm-forming capacity, revealing that not only does the capacity to form biofilms vary, but that variation also exists with respect to the molecular signals that initiate the response. Furthermore, when we tested three additional strains of *B. theta* we observed that LCA caused a substantial increase in biofilm formation for *Bt*-VPI-5482, while it had little impact on *Bt*-0940-1. Thus, significant strain-level variation exists with respect to the bile acid species to which individual strains are responsive to, suggesting that they show distinct biofilm responses depending on the biochemical milieu in which they reside. Additionally, *B. thetaiotaomicron* and related *Bacteroides* are known to have bile salt hydrolases (BSH) that are able to unconjugate secondary bile acids and/or modify these into tertiary bile acids (82–84). Therefore, although our results are clear that TLCA and GLCA are able to cause a profound increase in biofilm formation, we cannot exclude that actions of these BSH enzymes are modifying GLCA and TLCA into other forms. These results are interesting in the context of intestinal diseases as bile acids are known to have large fluctuations in patients with inflammatory bowel disease (IBD) predominately in patients with active disease including diarrea where the pool of secondary bile acids is largely diminished, while the pool of primary bile acids increases (85, 86). Without these secondary bile acids, it is possible that members of the gut microbiota such as *Bacteroides* are lacking important molecular cues which may dictate survival patterns within the intestine. Additionally, recent papers suggest that the repertoire of secondary bile acids and in-turn conjugated secondary bile salts has been understated due to the observance that BSH from the gut microbiome can produce amidated-bile acids through conjugating additional amino acids beyond taurine and glycine (84). These *bsh* gene functions are also important in shaping the gut microbiome and restricting the growth of pathogens like *Vibrio cholerae* (87) and *C. difficile* (73), including in a lithocholic acid-dependent mechanism (88). Beyond bile acids acting as one gut-derived molecule affecting biofilm formation, there exists a plethora of additional molecules and situational-dependency whereby substances such as short-chain fatty acids (89, 90) and carbohydrates affect biofilm formation of gut microbiome members (91, 92).

Interestingly, in testing individual bile acids, LCA and its related forms were consistently the only bile acid substrate capable of elciting or enhancing biofilm formation of the bile acid species we tested. We observed that non-LCA bile acids suppressed or had no impact on biofilm formation for intrinsic biofilm forming strains, *Bt-*0940-1 and *Bt*-5951. Further, for *Bt*-VPI-5482, no individual bile acid besides LCA and its related forms were capable of stimulating biofilm formation. Notably, bile is known to stimulate biofilm formation in these two strains and our data show that LCA concentrations are far below those supporting of biofilm formation, which suggests that (i) bile-induced biofilms may require more than one bile acid or a net sum of inducing bile acids or (ii). the non-bile acid fraction of bile, such as retinoic acid (22) or bilirubin (25) may promote a pro-biofilm state. Importantly, in our analysis of bile, no individual LCA compound or the sum of LCA compounds were at levels seen to stimulate biofilm formation. Lastly, in terms of substrates that play a role in the lifecycle of biofilm formation, our work demonstrates that the short-chain fatty acid acetic acid, is able to abolish the intrinsic biofilm forming propensity of strains *Bt-*0940-1 and *Bt*-5951 at concentrations that are physiologiocally relevant. SCFAs have previously been associated with inhibiting biofilm formation for pathogens (90, 93, 94) however, little is known about their role in biofilms of gut commensals. Given that *Bacteroides* are producers of SCFAs through the breakdown of carbohydrates which primarily leads to the production of acetate (4); it stands to reason that strains may have adapted a positive feedback mechanism whereby they sense acetate levels as a proxy for cell behavior based on nutrient availability in a similar manner as having been seen for bile acid concentrations affecting the gene regulation of carbohydrate metabolism (82). Therefore the ability to sense differences in both the bile acid pool and the SCFA pool during disease states may act as a molecular switch for *Bacteroides thetaiotaomicron,* signaling when it should exist in a biofilm for survival, versus as a planktonic cell when conditions are favorable.

Collectively, our work provides novel insights into the mechanisms through which *B. thetaiotaomicron* biofilm formation is mediated and uncovers strain-level variation not only in the capacity to form biofilms, but additionally with respect to the signals that coordinate this process. Given that we observed difference at the strain level, our approach underscrores the importance of examining physiological responses at the strain level to gain better understanding into the complex interplay of the inhabitants of the human gut and not being solely reliant on species level inferences as is common in DNA sequencing based approaches to study the microbiome. These strain-level variations may represent a distinct strategy through which individual strains promote their own fitness in the gut, or be reflective of their lifestyle in the intestine, and requires further investigation.

## Materials and Methods

### Bacterial Strains and Growth Conditions

All bacterial strains used in this study are listed in **Table S1**. *B. thetaiotaomicron* strains were routinely grown in tryptone-yeast extract-glucose (TYG) broth in an anaerobic chamber (Coy Manufacturing, Grass Lake, MI) at 37°C in an atmosphere of 10% CO_2_, 5% H_2_, balance N_2_. TYG was prepared as follows: Tryptone Peptone (10 g/L; Gibco, Cat. # 211921), Bacto Yeast Extract (5 g/L; BD, Cat. # 212750), Glucose (2 g/L; Sigma, Cat. # G8270), Cysteine HCl (0.5 g/L; Sigma, Cat. # C1276), 1M Potassium Phosphate Buffer (pH 7.2, 100 mL/L) made using 1M potassium phosphate monobasic (Sigma, Cat. # P0662), and 1M potassium phosphate dibasic (Sigma, Cat. # P3786), Vitamin K3 (menadione) (1 mL/L of a 1 mg/mL solution; Sigma, Cat. # M5625), TYG Salts (40 mL/L) made using MgSO4.7H2O (0.5 g/L; Sigma, Cat. # M63-500), NaHCO3 (10 g/L; Sigma, Cat. # S4019), and NaCl (2 g/L; Sigma, Cat. S3014), FeSO4 (1 mL/L of a 0.4 mg/mL stock; Sigma, Cat. # F8048), CaCl2 (1 mL/L of an 8 mg/mL stock; Sigma, Cat. # C7902), Hemin (0.5 mL/L of a 10 mg/mL solution; Sigma, Cat. # 51280), and Vitamin B12 (0.5mL/L of a 0.01mg/mL solution; Sigma, Cat. # V2876). For biofilm assays, bacteria were grown in either Brain Heart Infusion-supplemented broth with hemin and L-cysteine (BHIS) (95) or TYG broth where indicated. All liquid broth media was filter sterilized through 0.22µm filter (Millipore). Media formulations and supplier information for the above-mentioned media can be found in **Table S1**. For maintenance of *B. thetaiotaomicron* on solid media, Brain-Heart Infusion (BHI) plates supplemented with 10% v/v defibrinated horse blood (QuadFive, MT; Cat. # 210-500ml) were used (“BHI-blood plates” hereafter).

### Colony forming unit (CFUs) assay

Enumeration of CFUs was performed by spot-dilution plating on BHI-blood plates following biofilm assays after 48 hours of growth in 96-well plates. Briefly, 10 µL of culture was serially diluted and then 10 µL of these dilutions were spot plated onto plates in technical duplicate and allowed to incubate for 48 hours anaerobically followed by visual counting of colonies.

### 96-well biofilm assay

*B. thetaiotaomicron* strains were grown overnight in TYG broth and subcultured into either fresh TYG or BHIS with a wash step in fresh media via centrifugation at 8000 x g for three minutes and resuspended in the appropriate media at an OD_600_ of 0.01. Bacteria were transferred to a 96-well, non-treated polystyrene plate with round wells and flat bottom (Corning, Cat. # CLS3370), and grown in 200µL of media for 48 hours as indicated without disturbing the plate. Outside-facing wells of the plate were filled with water and no bacteria were seeded in these wells due to evaporation concerns. Assays involving enzyme treatments of DNase I (Sigma Cat. # 10104159001), RNase A (Thermo Scientific Cat. # J61996) (96), and Proteinase K (Fisher Scientific Cat. # BP1700) (68) were done via addition of these enzymes at the time of bacterial seeding into the 96-well plates and were left active for the duration of the 48 hour assay at the following concentration DNase I, 100U/ml, RNase A, 1U/ml, Proteinase K, 1mg/ml. Biofilm staining was performed as described previously (19). Briefly, bacterial supernatant was carefully removed from wells via pipette aspiration, followed by immediate fixing through the addition of 150µL Bouin’s solution (Sigma, Cat. # HT10132). Fixation was allowed to proceed for 20 minutes, following which the fixative was aspirated and the wells were subjected to gentle washing with 200µL ddH_2_O, with a total of three washes performed. 150µL of a 1% w/v Crystal Violet solution (Sigma, Cat. # V5265) was added and stained for 20 minutes followed by three, 200µL water washes. Stained biofilms were dissolved in a 4:1 ethanol:acetone mixture and were mixed via pipetting until all solids had dissolved. Absorbance was read at 575 nm on a microplate reader (Synergy HT, Biotek/Agilent Systems) using Gen 5 software. For those wells whose *A*_575_ exceeded the linear range of the plate reader, a dilution of the well contents was prepared for *A*_575_ measurement, and the true *A*_575_ calculated by correcting for the dilution factor.

### Testing the impact of bile acids and short-chain fatty acids on biofilm formation

The impact of bile acids on biofilm formation in TYG media was assayed via addition of the following compounds: sodium cholate hydrate (Sigma, Cat. # C9282) or conjugated bile acids were dissolved in ddH_2_O and diluted in TYG at the indicated molarity in each assay, and the following conjugated bile acids were used: glycocholic acid hydrate (Sigma, Cat. # G2878), sodium glycochenodeoxycholate (Sigma Cat. # G0759), sodium glycocholate hydrate (Sigma, Cat. # G7132), taurocholic acid sodium salt hydrate (Sigma, Cat. # T4009), sodium taurochenodeoxycholate (Sigma, Cat. # T6260), and sodium taurodeoxycholate hydrate (Sigma, Cat. # T0875). For bile acids not readily soluble in water, 100% ethanol or 100% DMSO was used to make stock solutions at high molar concentrations including lithocholic acid (Sigma Cat. # L6250), glycolithocholic acid (Cayman Chemical Cat. # 20273), taurolithocholic acid (Cayman Chemical Cat. # 17275), alloisolithocholic acid (Cayman Chemical Cat. # 29542), isolithocholic acid (Cayman Chemical Cat. # 29545), dehydrolithocholic acid (also known as 3-oxo-LCA; Cayman Chemical Cat. # 29544), deoxycholic acid (Sigma Cat. # D2510), hyodeoxycholic acid (Sigma, Cat. # H3838), ursodeoxycholic acid (Sigma, Cat. # U5127), and chenodeoxycholic acid (Sigma, Cat. # C9377). These solutions were then diluted in TYG to yield working solutions containing 2% v/v ethanol or 1% v/v DMSO at the highest concentration of these compounds, the compounds were than mixed 1:1 in the well with 2x concentration of TYG with bacteria, yielding a final concentration of 1% or 0.5% v/v ethanol or DMSO. To control for ethanol or DMSO affecting biofilm formation, TYG media containing 2% v/v ethanol or 1% v/v DMSO without added bile salts were used as controls, and comparisons for individual bile acids were made to the appropriate controls. Dilutions were made fresh in all cases and were diluted in the well as addition of the stock into TYG caused some of the compound to precipitate. To control for background absorbance caused from this precipitation, blank wells without bacteria were run as controls and stained as above. Biofilm assays were grown for 48 hours prior to staining. As a control for biofilm formation, bile (Sigma Cat. # B8381) was used as described previously for *B. thetaiotaomicron* strains (18). Briefly, bile was added as a percent w/v into liquid TYG media at 0.0625, 0.125, 0.25, 0.5, 1, or 2 % w/v to test the response of several isolates of *B. thetaiotaomicron*. For all bulk bile, bacteria were grown for 48 hours and then a crystal violet-based assay to measure biofilm formation was performed. For short-chain fatty acid (SCFA) biofilm assays, the compounds were added to TYG as above for bile acids, followed by addition of bacteria and allowed to incubate undisturbed for 48 hours followed by staining with Crystal Violet as above. Acids were commercially purchased from the following vendors and used at the indicated concentrations within each assay; acetic acid (Sigma, Cat. # A6283), propionic acid (Sigma, Cat. # 402907), isobutyric acid (Sigma, Cat. # 58360), isovaleric acid (Sigma, Cat. # 129542), and valeric acid (Sigma Cat. # 240370).

### Bile acids quantification

Bile acids were quantified by a previously published a stable-isotope-dilution liquid chromatography mass spectrometry (LC-MS/MS) method (97) with some modifications. Briefly, ice-cold methanolic internal standard solution (80 µl) containing D_4_-chenodeoxycholic acid (D_4_-CDCA), D_4_-cholic acid (D_4_-CA), D_4_-deoxycholic acid (D_4_-DCA), D_4_-lithocholic acid (D_4_-LCA), D_4_-glycochenodeoxycholic acid (D_4_-GCDCA), D_4_-glycocholic acid (D_4_-GCA), D_4_-glycodeoxycholic acid (D_4_-GDCA), D_4_-glycolithocholic acid (D_4_-GLCA), D_4_-glycoursodeoxycholic acid (D_4_-GUDCA), D_4_-taurocholic acid (D_4_-TCA), D_4_-taurodeoxycholic acid (D_4_-TDCA) and D_4_-taurochenodeoxycholic acid (D_4_-TCDCA) was mixed with TYG media or TYG media plus bile at 1% w/v (20 µl), vortexed and centrifuged (14,000x*g*, 20 min, 4°C). To cover different concentration ranges media was run as undiluted as well as diluted 10x and 100x. Supernatant was transferred to glass HPLC vials with micro-inserts and subjected to LC-MS/MS analysis. Calibration standards were prepared in methanol and quality control samples were prepared from pulled plasma samples and processed identically as samples. LC-MS/MS analysis was performed on the same chromatographic system as described above. An ACE Excel C18-Amide column (75 mm × 2.1 mm; 1.7 μm) (Cat. # EXL-1712-7502U, Avantor, Radnor Township, PA) was used for chromatographic separation. A gradient of solvent A (0.1% acetic acid in water) and B (0.1% acetic acid in acetonitrile: methanol 50:50; v/v) was used for chromatographic separation with a flow rate of 0.4 mL/min and a 3 µL injection volume. Electrospray ionization in negative ion mode was used with the following multiple reaction monitoring (MRM) conditions: *m/z* 373.3→373.3 for cholenic acid and 3-keto-cholanic acid; *m/z* 375.3→375.0 for iso-LCA, LCA, allo-iso-LCA, and allo-LCA; *m/z* 377.3→377.3 nor-DCA; *m/z* 387.3→387.3 for 3,7-diketocholanic acid, 3,6-diketocholanic acid and 12-keto-9,5-cholenic acid; *m/z* 389.3→389.3 for 3-keto-DCA, 6-keto-LCA, 7-keto-LCA, 12-keto-LCA, apo-CA, 3-keto-CDCA; *m/z* 391.3→391.3 for DCA, hyodeoxycholic acid (HDCA), CDCA, UDCA, iso-UDCA, 7-iso-DCA, iso-DCA and muro-DCA; *m/z* 401.2→331.1 for triketocholanic acid; *m/z* 403.3→403.3 for 7,12-diketo-LCA, 7-keto-DCA and takeda ketol; *m/z* 407.2→407.2 for CA, α-muricholic acid (α-MCA), β-MCA, ω-MCA, hyocholic acid (HCA), ursocholic acid (UCA) and allo-CA; *m/z* 432.3→73.9 for GLCA; *m/z* 448.3→73.9 for GCDCA, GHDCA, GUDCA and GDCA; *m/z* 455.2→96.9 for LCA-SO4; *m/z* 458.2→73.9 for G-dehydroCA; *m/z* 464.3→73.9 for GCA, GHCA; *m/z* 482.2→123.9 for TLCA; *m/z* 487.2→96.9 for CA-7-SO4; *m/z* 498.2→123.9 for THDCA, TCDCA, TDCA and TUDCA; *m/z* 514.2→123.9 for TCA, THCA, T-α-MCA, T-β-MCA and T-ω-MCA; *m/z* 379.3→379.3 for D_4_-LCA; *m/z* 395.3→395.3 for D_4_-CDCA and D_4_-DCA; *m/z* 411.2→411.2 for D_4_-CA; *m/z* 436.2→73.9 for D_4_-GLCA; *m/z* 452.3→73.9 for D_4_-GCDCA, D_4_-GDCA and D_4_-GUDCA; *m/z* 468.2→73.9 for D_4_-GCA; *m/z* 502.2→127.9 for D_4_-TDCA and D_4_-TCDCA; *m/z* 518.2→127.9 for D_4_-TCA

### Statistical Analysis

Statistical analyses were performed using built-in analysis within GraphPad Prism version 10.2.1. The specific statistical tests used are indicated within the figure legends.

## Author Contributions

R.W.P.G. conceived of and performed all experiments involving biofilms, growth assays, and quantitative PCR, and was primarily responsible for writing the manuscript. M.J.E. and J.M.T. assisted in experiments and provided intellectual input. N.S. performed genome sequencing analysis of strains used. A.K. helped in design and optimization of biofilm experiments. I.N. performed assessments of bile in growth media. P.P.A. designed experiments, oversaw data analysis, and helped in writing the manuscript.

## Acknowledgments

The authors are indebted to Dr. Jeffrey I. Gordon and Dr. Janaki Lelwala-Guruge for the provision of *B. thetaiotaomicron* isolates used in these studies, and to Dr. Abigail A. Salyers for the original isolation of some of these strains.

---

**Figure S1.**
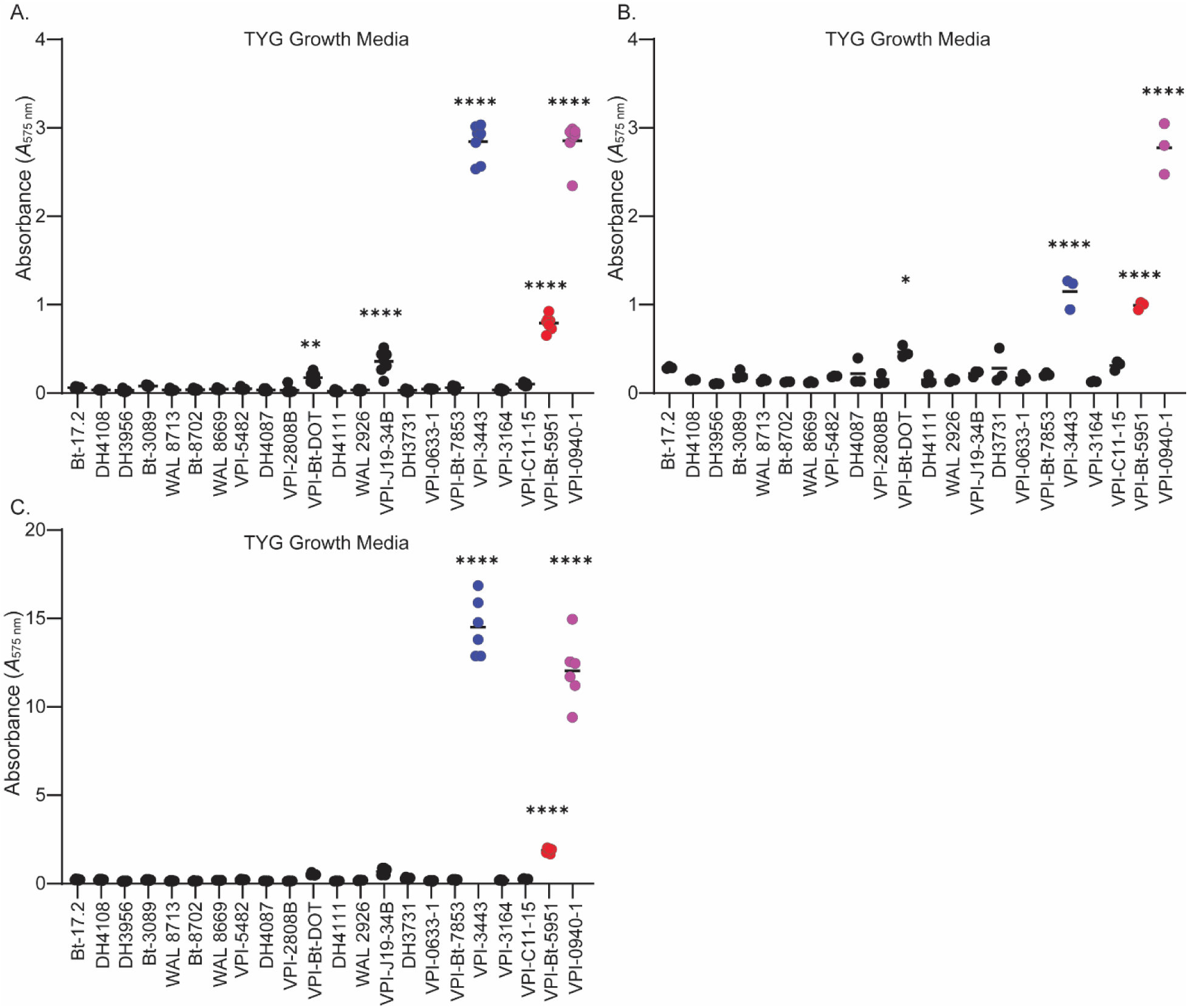
Replicates of biofilm formation in TYG media, related to Figure 1. (A-C) Independent replicates for the validation of the biofilm assay in TYG media, reflecting the natural variation that occurs in the magnitude of biofilm form, but reflective of the maintenance of the same pattern between individual strains. All biofilms were quantified at 48 hours post inoculation using a crystal violet assay. Statistical significance was determined using one-way ANOVA with Dunnet’s multiple comparisons to the mean of strain VPI-5482. *p* values of <0.05 were considered statistically significant, **, *p*<0.01 and ****, *p*<0.0001 for all graphs shown.

**Figure S2.**
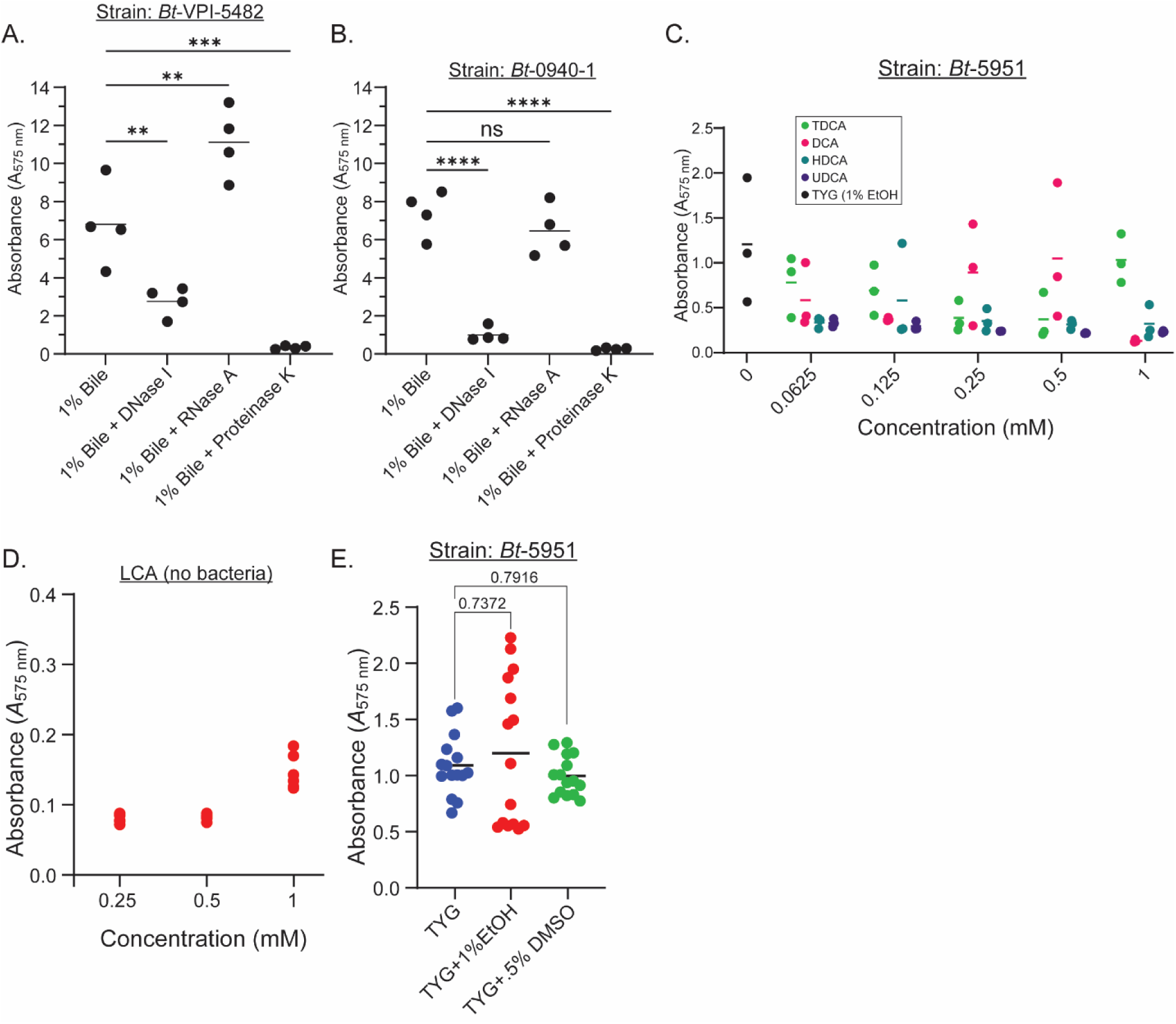
Impact of bile on biofilm formation, related to Figure 2. (A, B) Biofilm formation in TYG media with 1% w/v bile after 48 hours of growth for strains *Bt*-VPI-5482 (A) and *Bt*-0940-1 (B). In (A, B), DNase I, RNase A, and Proteinase K enzymes were added to assay biofilm formation induced by bile in the presence of these enzymes. Graphs show technical replicates with a line at the mean of the data and graphs are representative of 2 independent experiments. (C) Biofilm formation of strain *Bt*-5951 at 48 hours in TYG media containing doses of hyodeoxycholic acid (HDCA), ursodeoxycholic acid (UDCA), taurodeoxycholic acid (TDCA), and deoxycholic acid (DCA). Data is representative of 2 independent experiments. (D) Biofilm assay of lithocholic acid (LCA) added to TYG media without added bacteria at the indicated concentrations. Data is representative of 2 independent experiments. (E) Biofilm assay of strain *Bt-*5951 grown in TYG or TYG in the presence of 1% (w/v) ethanol (EtOH) or 0.5% DMSO (w/v). Statistical significance was determined using a one-way ANOVA with Dunnet’s multiple comparisons to 1% bile in (A, B), or compared to TYG without EtOH and DMSO added (E). *p* values of <0.05 were considered statistically significant **, *p*<0.01 and *\*\*\*, p*<0.001 and ****, *p*<0.0001.

**Figure S3.**
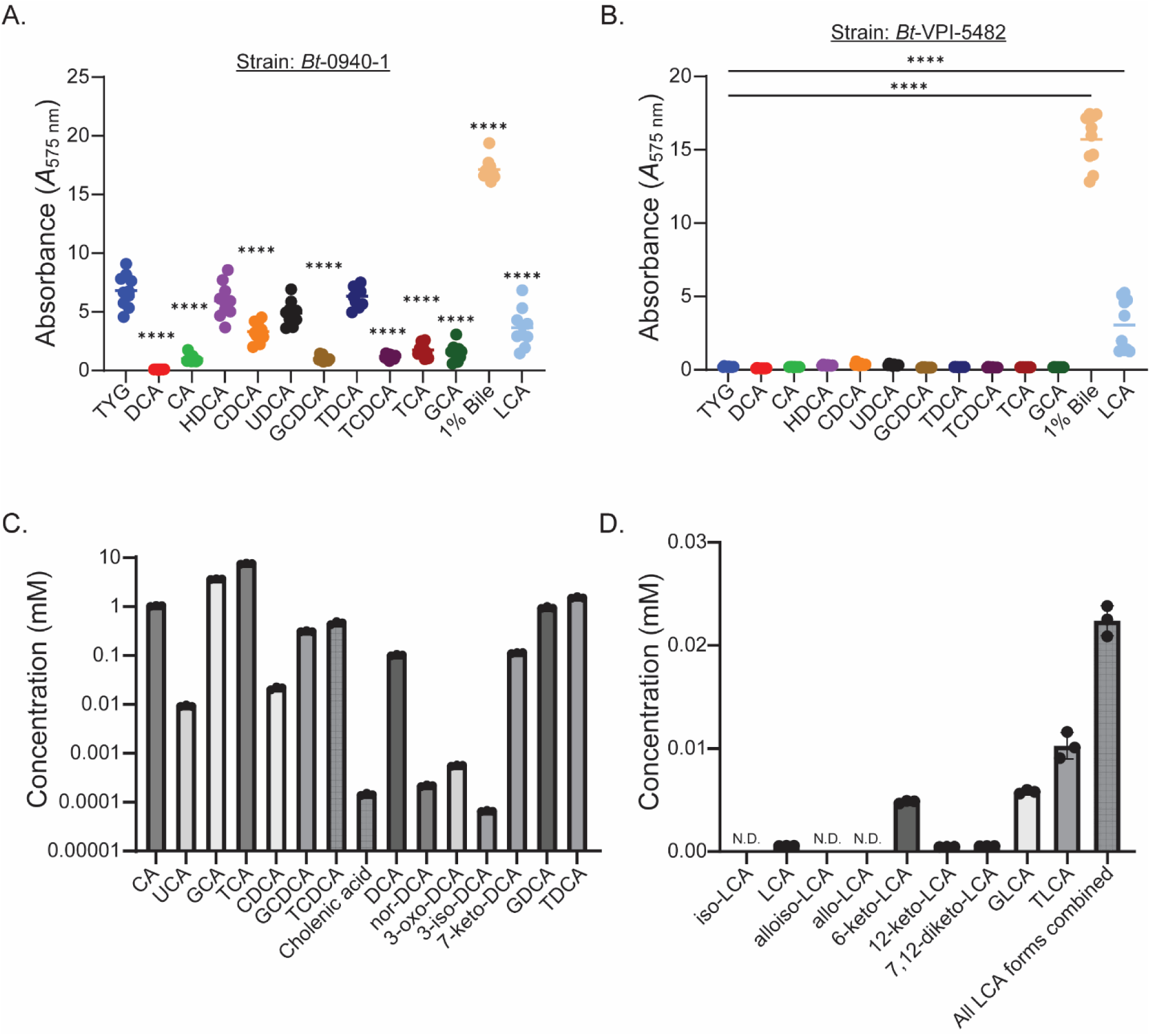
Lack of non-LCA bile-acid induced biofilm formation and compositional analysis of bile, related to Figure 3. (A,B) Biofilm formation for strain *Bt-*0940-1 (A) or *Bt*-VPI-5482 (B) grown in TYG for 48 hours with bile acids added at 0.5 mM or bile at 1% (w/v). All bile acids were significantly lower than TYG, except hyodeoxycholic acid (HDCA) and taurodeoxycholic acid (TDCA) in (A), significance is indicated by asterisks above the treatments significant compared to TYG. Other bile acids tested include DCA=deoxycholic acid, CA=cholic acid, acid, CDCA=chenodeoxycholic acid, UDCA=ursodeoxycholic acid, GCDCA=glycochenodexocycholic acid, TCDCA=taurochenodexocycholic acid, TCA=taurocholic acid, GCA=glycocholic acid, and LCA= lithocholic acid. (B,C) Biofilm formation in TYG media supplemented with the indicated bile acid at 0.5 mM for all substrates or 1% bile (w/v). Data is representative of 3 independent biological replicates. (C,D) Mass spectrometry analysis of 1% (w/v) commercially available bile used to induce biofilm formation. Data shown are 3 technical replicates showing bile acid analyates detected above a calibrated limit of detection. Both (C) and (D) are from the same sample of bile, but have been separated here for ease of viewing into non-LCA bile acids (C) and LCA bile acids (D) as the concentration of LCA bile acids are orders of magnitude lower. In (D), we are showing a bar at the far right of the graph that has pooled all LCA forms displayed on this panel to show the total sum of all LCA epimers and conjugated forms. Statistical significance was determined using a one-way ANOVA with Dunnet’s multiple comparisons to TYG in (A, B), *p* values of <0.05 were considered statistically significant ****, *p*<0.0001.

**Figure S4.**
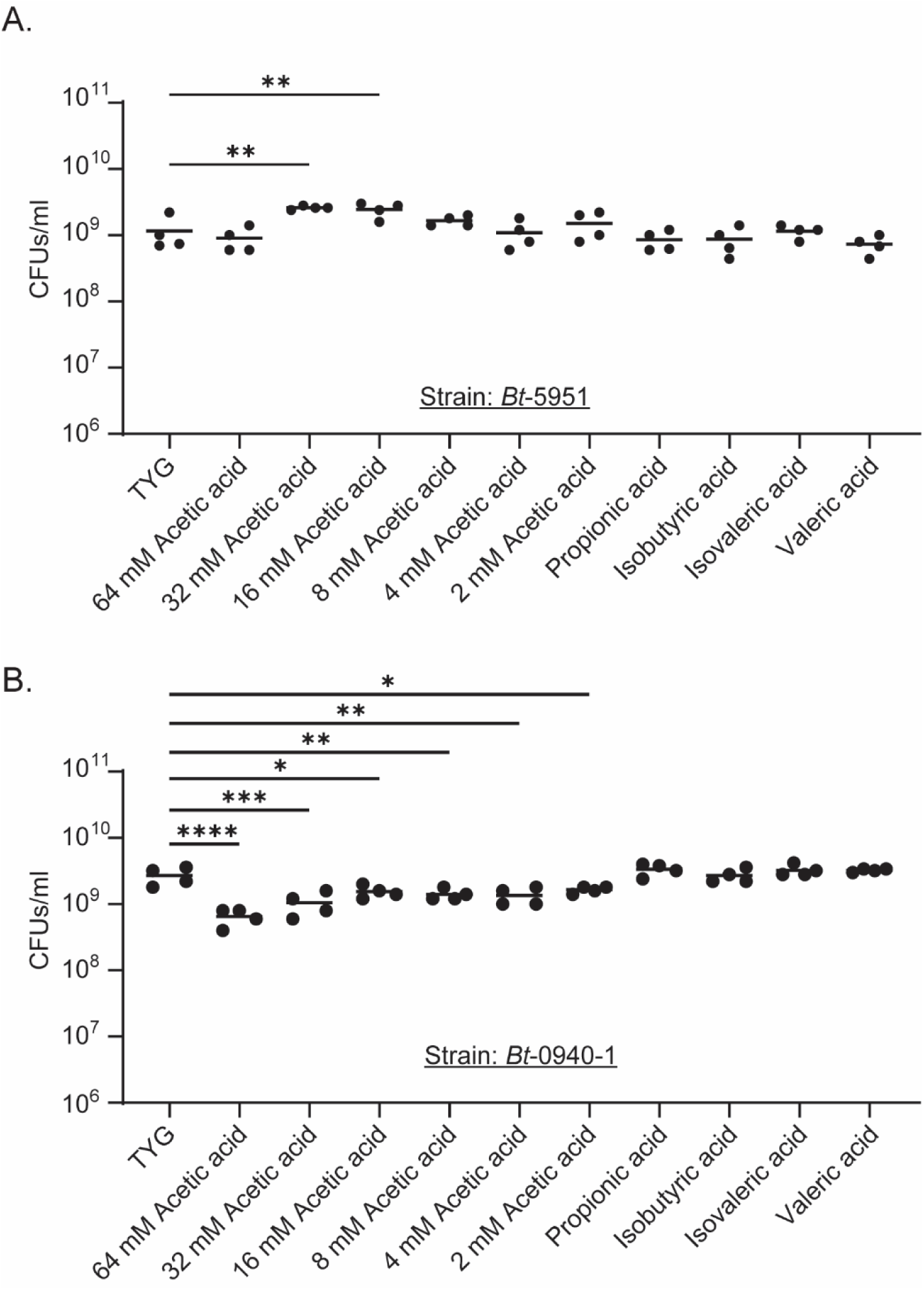
Short-chain fatty acids do not substantially affect viability, related to Figure 4. (A-B) Colony forming unit (CFU) assay with n=4 biological replicates run concurrently for strains *Bt*-5951 (A) and *Bt*-0940-1 (B). Bacteria were plated on BHI-blood agar after 48 hours of growth in TYG media with the indicated concentration of acetic acid added or propionic acid (8 mM), isobutyric acid (4 mM), isovaleric acid (1 mM), or valeric acid (1 mM). Data is representative of 2 independent experiments. Statistical significance was determined using a one-way ANOVA with Dunnet’s multiple comparisons to TYG without acetate added (0 mM), *p* values of <0.05 were considered statistically significant, *, *p*<0.05, **, *p*<0.01, *\*\*\*, p*<0.001, and ****, *p*<0.0001.

